# A geometrical model of cell fate specification in the mouse blastocyst

**DOI:** 10.1101/2023.10.23.563683

**Authors:** Archishman Raju, Eric D. Siggia

**Affiliations:** Simons Centre for the Study of Living Machines, National Centre for Biological Sciences, Tata Institute of Fundamental Research, Bangalore 560065, India; Center for Studies in Physics and Biology, Rockefeller University, New York, NY 10065, USA

## Abstract

The lineage decision that generates the epiblast and primitive endoderm from the inner cell mass (ICM) is a paradigm for cell fate specification. Recent mathematics has formalized Waddington’s landscape metaphor and proven that lineage decisions in detailed gene network models must conform to a small list of low dimensional stereotypic changes called bifurcations. The most plausible bifurcation for the ICM is the so-called heteroclinic flip that we define and elaborate here. Our reanalysis of recent data suggests that there is sufficient cell movement in the ICM so the FGF signal, which drives the lineage decision, can be treated as spatially uniform. We thus extend the bifurcation model for a single cell to the entire ICM by means of a self-consistently defined time-dependent FGF signal. This model is consistent with available data and we propose additional dynamic experiments to test it further. This demonstrates that simplified, quantitative, and intuitively transparent descriptions are possible when attention is shifted from specific genes to lineages. The flip bifurcation is a very plausible model for any situation where the embryo needs control over the relative proportions of two fates by a morphogen feedback.

## 1 Introduction

One of the hallmarks of development is the robust allocation of fates which is tolerant both to noise and external insults. This is often represented as Waddington’s metaphor of flow down a landscape where lineage decisions correspond to valleys splitting [1]. This metaphor has recently been formalized into a mathematical tool for constructing landscapes and fitting cell fate data [2].

The pre-implantation mouse blastocyst provides an ideal example of robust allocation of fates, because it is experimentally amenable to ex-vivo manipulation, highly reproducible, and extensively studied [3]. The early development of the mouse embryo is relatively well understood and takes place through two binary decisions [4]. First, the blastomeres differentiate into Inner Cell Mass (ICM) and Trophectoderm (TE). This is followed by the differentiation of ICM into Epiblast (Epi) or Primitive Endoderm (PrE) which happens between E3.25 to E4. The Epiblast becomes the embryo proper with the PrE making supporting tissue. The FGF signaling pathway is known to regulate the second of these two transitions though the exact mechanism through which it acts is still under debate [5, 6]. Two receptors respond to the FGF ligand, FGFR1 is present in all ICM cells, and FGFR2 in PrE only, but both contribute to the lineage decision and partially compensate [7,8]. The receptors activate a downstream MAP kinase pathway and ERK signaling. Cell fates are usually tracked through the activity of two transcription factors: NANOG, required for specification of Epi and GATA6, required for specification of PrE [9, 10].

Our present study is motivated by a set of recent experiments which measured the dynamics of ERK levels through a Kinase Translocation Reporter (KTR). The reporter acts as a substrate for the ERK and localizes from the nucleus to the cytoplasm when ERK is activated. Thus the cytoplasm to nuclear (C:N) ratio of the reporter acts as a proxy for the ERK activity. This in turn, is a proxy for the level of the FGF signal. The experiments by Simon et al. [11] took two sets of measurements. First, they measured the ERK activity for a 2 hour period at intervals of 5 minutes during the critical period for fate specification between E3.25 to E3.75. They observed a considerable amount of heterogeneity in the ERK levels but found mean ERK levels over the 2 hour period to be substantially higher in the prospective PrE cells than the Epi cells by staining fixed cells for NANOG and GATA6 at the end of the live reporter measurement. Second, to check the consistency of their short term results, they took some long-term measurements of ERK activity in the early blastocyst (∼ E3.25) for a 12 hour period with a 15 minute interval for measurement. (A contemporaneous study [12], with no direct relevance for us, found a correlation between the ERK activity in the hour following mitotic exit, and lineage selection.)

Previous theoretical studies have mathematically modeled the blastocyst using a spatially localized range of FGF signaling and a model of the gene regulatory networks. The most extensive of these models is by Tosenberger et al. [13], which proposes a gene regulatory circuit with tristability and is consistent with available experimental data perturbing FGF levels by adding it to the media or drug inhibition [14]. Another study modeled the ICM as a bistable system, and compared their mathematical model to experiments which showed that the embryo could maintain a robust composition of cells even when lineage-restricted cells were added or existing specified cells were ablated *in-vivo* [15]. Another recent model proposed a collective transition where bistability in the system is the result of a pitchfork bifurcation governed by the number of cells [16].

We propose a compact ‘geometric’ model for this transition following Ref. [2]. We reduce the number of variables to the mathematical minimum, show some prior models relied on parameter tuning, and expose the common features of the ICM transition to other examples of the flip bifurcation. We rigorously extend the landscape metaphor, generally construed as a single cell, to the entire ICM. The key is a reanalysis of the data in Ref. [11], that argues that cells in the ICM are sufficiently mobile so they see a spatially averaged, but time dependent, FGF level.

Geometric models exploit the parallel phenomenology between experimental embryology and dynamical systems theory. Cell fate specification relies on a gene regulatory network which can be modeled as a set of differential equations. Motion in that space can be thought of as a *flow*. Stable fixed points (henceforth just *fixed points*) attract all near by points. The fixed points are where the flow stops (or is committed to a valley whose subsequent development is not followed). Fixed points correspond to cell types at the relevant time-scales. Saddle points, which have both stable and unstable directions, are where the cell makes a decision. Geometrical models focus on the topology of the flows between the saddles and fixed points [2, 17–20]. They model the dynamics in a low-dimensional abstract space which is to be understood as capturing the important part of the full gene-expression dynamics. Crucially, cell fates are modeled but not the genes themselves.

We consider two geometrical models for how the transition from ICM to Epi or PrE could take place. We compare the two models to elucidate the difference in their predictions for this fate decision. We show how our preferred geometrical model is consistent with existing experimental data of FGF perturbations and the robust allocation of cell fates in the face of insults. Finally, we discuss how more controlled dynamic experiments can decisively distinguish between alternate geometries.

## 2 Results

### 2.1 Analysis of Spatial Correlations in ERK activity

We first analyzed the live imaging data of ERK activity from Ref. [11] to measure spatial correlations in the ERK activity, as a proxy for the FGF. If there are very local cell-cell interactions mediated by the FGF, then we expect the presence of strong spatial correlations in the ERK signal. Alternatively, if cell movement and diffusion of FGF homogenizes the FGF concentrations, then we expect very little spatial correlation in ERK activity.

The first fate decision is between ICM and TE cells. These two sets of cells are spatially separated and manually labeled in the data. We remove the data corresponding to cells that have already specified to TE and focus only on the ERK activity in the ICM cells.

To estimate the affect of the cell movement, we consider a Voronoi tessellation of the cells based on a nuclear marker. We then find the unique number of neighbors lost and gained during the 2 hr interval of the measurement. We show a sample embryo with cell movement in Figure 1D. The movement of a few typical cells in the embryo is shown alongside the minimum bounding ellipsoid. The cell movement covers a significant fraction of the available space after the blastocyst cavitates. We estimate that a neighbor is lost or gained (in the Voronoi sense) on average every 25 minutes (see Materials and Methods).

**Figure 1:**
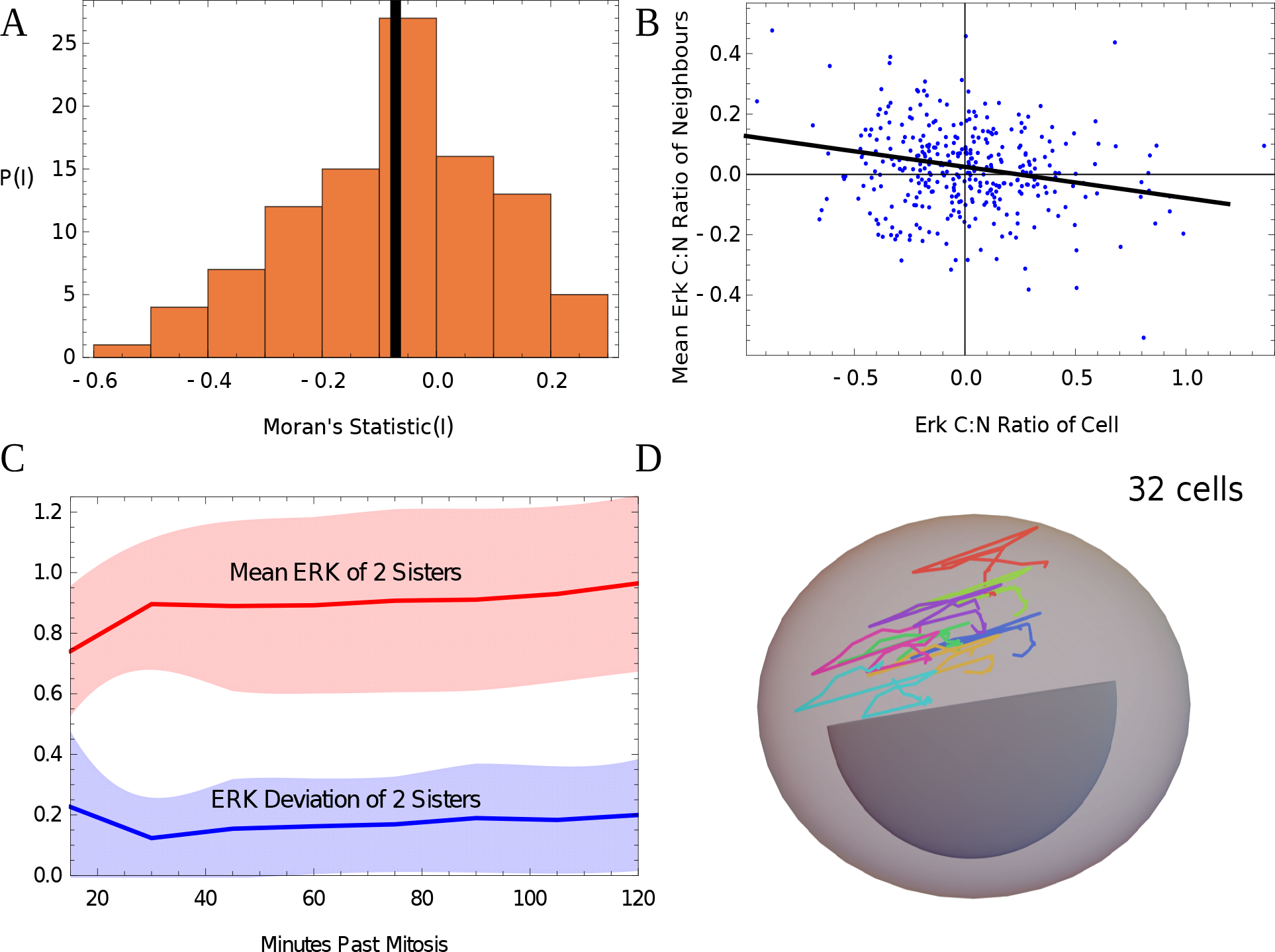
Analysis of correlations of live imaging data of ERK activity. These figures examine the data in Ref. [11].(A) A histogram of the Moran’s I index, which is a measure of spatial correlations, with -1 for fully anti-correlated and 1 for fully correlated data. The index is calculated by looking at snapshots separated by 30 minutes. The ERK activity (C:N ratio) of every cell in 25 embryos is correlated with that of its nearby cells (at a distance of 20 *μ*m). A small negative mean correlation (solid black line) is obtained equal to − 0.07. This is significantly different from 0 (p value 0.0001) but small compared to −1. (B) A plot of the ERK activity alongside the average value for its neighbors by taking one random snapshot from all 25 embryos. Each embryo has an average of about 14 cells (excluding TE cells) for a total of 346 cells used in the plot. The best fit line is has a slope of − 0.1 reflecting small but non-zero anti-correlation. Note that the points shown are centered around the mean. (C) The mean ERK levels of two sister cells after mitosis plotted alongside the absolute deviation of ERK. The deviations are substantially smaller than the mean reflecting correlated lineages. A total of 111 cells are used for the plot with 8 data points each. Shaded regions are one standard deviation. (D) A representative set of trajectories of 8 cells from one embryo at the 32 cell stage is shown for the duration of 2 hours. The outer dome is the minimum bounding ellipsoid of all trajectories including the Trophectoderm (TE) cells which have already segregated. The region marked in dark gray roughly shows the position of the cavity. Different colors represent different cells.

We measure the spatial correlations in the data of Simon et al. by looking at snapshots of the embryos and then seeing how correlated the ERK activity is. To obtain independent samples, we only take snapshots which are separated by a time of 30 minutes in the same embryo, to account for the time-scale at which cells lose or gain a neighbor. We use the Moran’s I statistic to test the significance of the spatial correlations. A value of −1 of the statistic indicates perfect anti-correlation, a value of 1 indicates perfect correlation and 0 implies no correlation.

There are two plots drawn using the Moran’s I statistics which bring out the lack of significant spatial correlation in the ERK activity shown in Figure 1. The first is a histogram of the I values across the 25 different embryos (with 4 data points per embryo). The mean statistic is − 0.07 *±* 0.02. The statistic is significantly different from 0 with a p-value of .0001. This implies that there is some statistically significant negative spatial correlation but the mean is small implying the correlation is very weak. A second way to visualize this is to plot the value of normalized ERK activity alongside the average normalized value for the neighbors. This plot also indicates the presence of a weak albeit non-zero spatial negative correlation.

Next, we look at the lineage statistics in the 12 hour movies and show the deviation in the ERK activity of two sister cells right after division. The deviations are substantially smaller than the mean ERK levels of the two sisters as shown in Figure 1C. This reflects positive lineage correlations in agreement with Ref. [12], but since we find no substantial spatial correlation, cell-movement must scramble the position of daughter cells. We neglect the role of lineage correlation between sisters in the eventual end fate, for reasons explained in the Discussion.

These results all imply that FGF diffusion and cell movement together do not allow the persistence of any strong spatial correlations in ERK activity. Thus we model FGF as a spatially uniform but time dependent signal that acts equally on all cells. Its value is the sum of contributions from each cell as a function of its internal state, i.e., its position on the landscapes in Figure 2, whose specification is an important part of the model.

**Figure 2:**
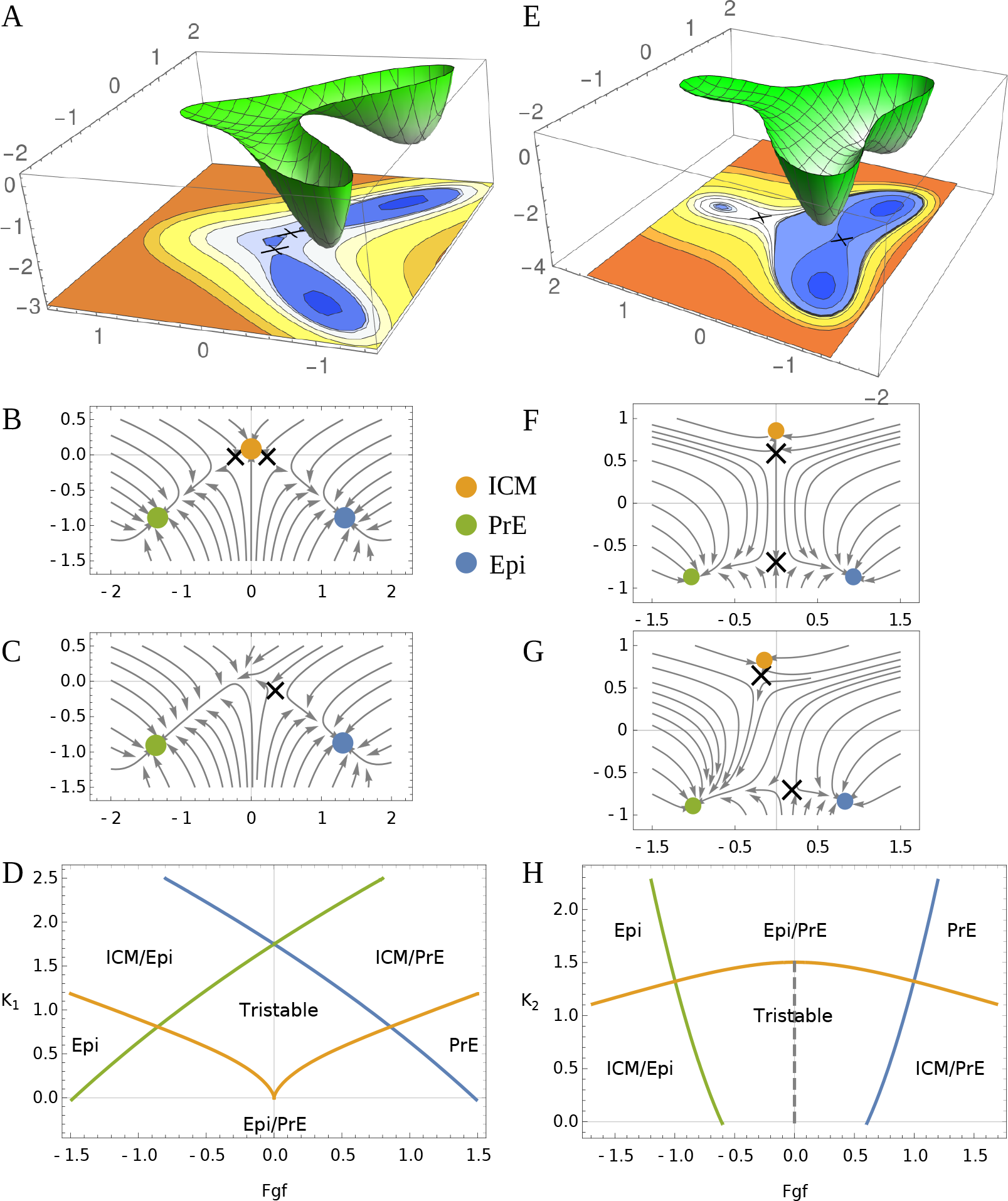
Two possible geometries for the ICM state near its bifurcation point. (A) The dual cusp geometry has a potential (schematic) where the central state (ICM) is connected to Epi and PrE but one can not directly transition from Epi to PrE without passing through the ICM. (B) Flow for the dual cusp *K*_1_ = 0.15 (C) If the FGF concentration is increased, a bifurcation of the ICM state occurs and the flow goes towards PrE (green). (D) The full bifurcation diagram for the dual cusp. Solid lines indicate points of bifurcation where one of the states disappears with the color indicating which state (*K*_1,2_ controls the tilt of the potential in the vertical direction). (E) A schematic potential for the heteroclinic flip where Epi and PrE are directly connected. (F) The flow for the heteroclinic flip with *K*_2_ = 1.4 with the FGF parameter set to the unstable point where the flow from the upper saddle hits the lower one. (G) If the FGF concentration is increased, flows from the saddle near the ICM state go towards PrE but no bifurcation occurs. (H) The full bifurcation diagram for the heteroclinic flip. Solid lines show the point at which bifurcations occur as in (D) and the dashed line indicates when the flow out of the upper saddle precisely hits the lower one. The bifurcation of the ICM state takes place at *K*_1_ = 0 for the cusp and *K*_2_ = 1.5 for the flip.

### 2.2 Constructing the geometrical model

Our recent work, [2], suggests two possible geometries for the ICM lineage decision that we model as a tristable dynamical system with the ICM, Epi, and PrE as the three available states. The two geometries are shown in Figure 2, as landscapes and corresponding flows downhill. The landscapes are described by two parameters, one controls the stability of the ICM state and is primarily a tilt in the vertical direction, while the second is the effect of FGF which tilts horizontally, Figure 2D,H.

The dual cusp, Figure 2A,B, is a local bifurcation in that the flow intersecting a small circle around the ICM and two saddle points denoted as X in Figure 2B is identical to that for a circle around the single saddle in Figure 2C. The dual cusp is a single *point* in the two dimensional parameter space at which an ICM cell looses stability towards Epi or PrE. The final outcomes may be biased between Epi and PrE but both are accessible. The dual cusp geometry is effectively one dimensional, all the decisions happen on the flow curve connecting the three states (thus vertical and horizontal tilts in the landscape may have similar effects).

In the alternative flip geometry, the ICM looses stability along a *line* in parameter space, Figure 2D vs H, and in this sense does not require parameter tuning, while the dual cusp does. The flip bifurcation is global and involves two ‘decisions’, the first (local) to destabilize the ICM and the second (global) whether to flow to PrE or Epi. Both are ‘typical’ in that they occur along a line in parameter space. As we elaborate below, the ICM cells first flow down a confining valley towards the lower saddle point, then the valley opens becoming a ridge and the cells flow off the ridge to either PrE or Epi. The lower saddle separating the PrE and Epi is unstable to the slightest amount of noise. The flip geometry is inherently two dimensional and cells tending towards Epi can be diverted to PrE by exogenous FGF without passing through the ICM.

Finally, it’s not understood whether the signal that destabilizes the ICM around E3.25 is a reflection of cell number or absolute time, and its molecular identity. However, in a geometric model, all that matters is that the previously stable ICM becomes unstable by a bifurcation with the upper saddle point.

### 2.3 Quantifying the effect of FGF on a single cell

The two geometries make very different predictions for response to a pulse of FGF just after the ICM state has bifurcated away (Figure 3). If the cell is tending towards Epi, then a pulse of FGF under the dual cusp geometry will force the cell to revert to ICM on its way to the PrE state. The meaning of lineage in this context is that no other paths between Epi and PrE exist. For the flip on the other hand a direct transition between the two states in possible, more so if the FGF applied before the wells representing the states are too deep.

**Figure 3:**
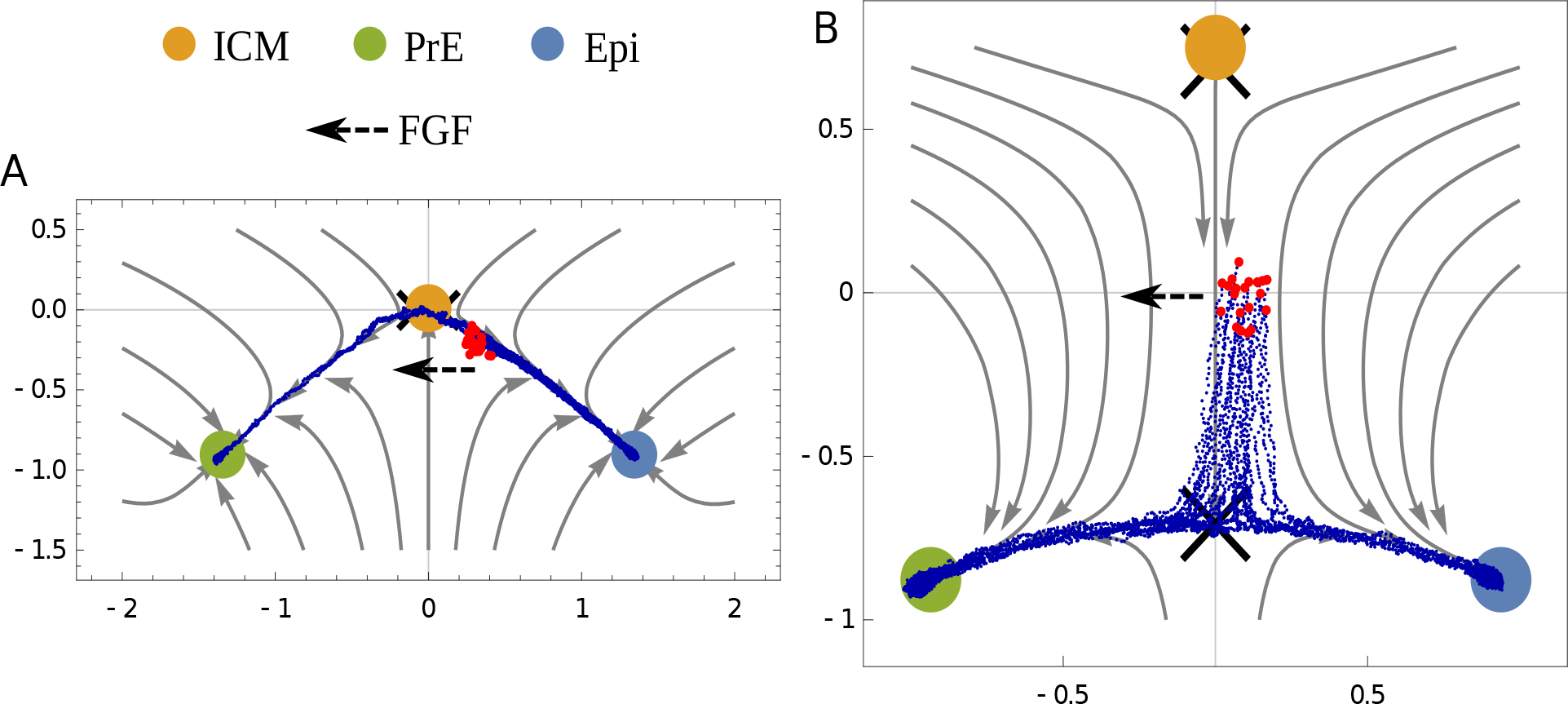
Transition of fate to PrE with FGF over-expression. To demonstrate the difference in the two geometries, we show how external FGF changes the fate of cells that are tending towards Epi but but not yet committed to that state near the time when the ICM state becomes unstable. Blue lines show trajectories for multiple cells whose initial conditions are shown in red. Dashed line shows the direction in which FGF acts. A fraction of the cells commit to PrE. (A) In the dual cusp geometry, going from Epi to PrE requires going through the ICM state. (B) In the heteroclinic flip, the cells can directly transition from Epi to PrE.

Timed perturbations would convey the most quantitative information about the structure of the landscape, and we consider again a landscape that is biased towards the Epi state and where the ICM state has just undergone a bifurcation. For the flip, as the cell exits the ICM state (Figure 4A), the flow is compressed and the state is brought towards the heteroclinic orbit connecting the two saddle points. This is akin to rolling down a valley. Thus an early pulse of FGF when the cell is in the valley will have little effect (Figure 4B) while a pulse when the cell is just exiting the valley will have maximal effect (Figure 4C,D). For the dual cusp the sensitivity decreases monotonically in time (Figure 4E).

**Figure 4:**
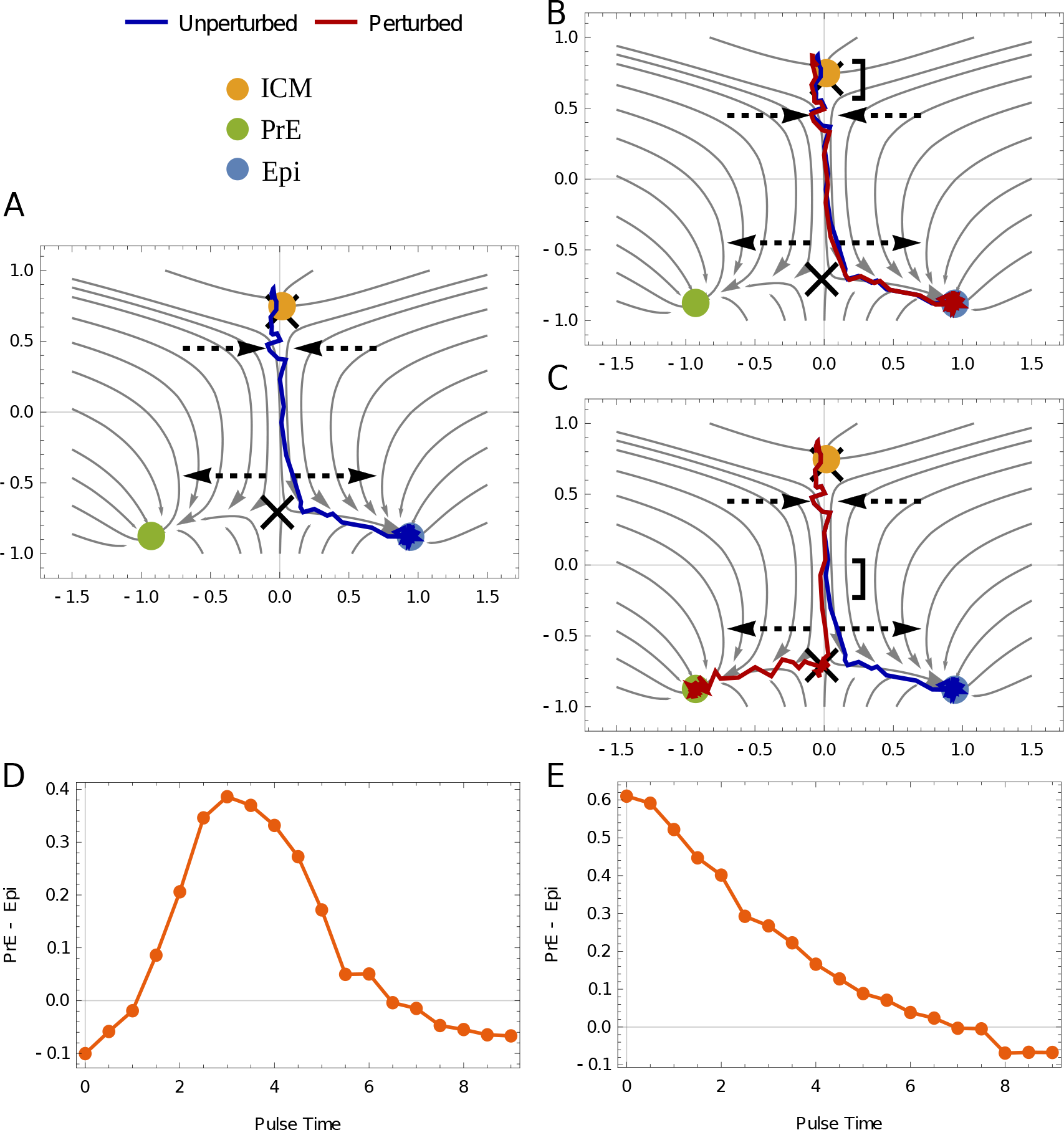
Time dependent perturbations differentiate the two geometries. A standardized pulse of FGF (magnitude 0.2, duration 0.5) is applied at various times to a topography slightly biased to Epi without internal FGF feedbacks to illustrate the two topologies. (A) A representative trajectory in the absence of an FGF pulse. Dashed arrows indicate regions where the flow converges and diverges, following Figure 2E. (B) If external FGF is put in early (square bracket), the trajectory (red) deviates slightly but has time to recover and moves back to Epi. (C) If FGF is put in late, the diverging flow amplifies the perturbation and the trajectory (red) moves towards PrE. (D) Sensitivity to timed FGF perturbations for the heteroclinic flip calculated as the fraction difference between PrE and Epi. (On this scale the WT embryo with 60% PrE would appear as 0.2 (= 0.6-0.4)). The limits at start and end times indicate the bias towards Epi (modeled as a constant negative magnitude 0.004 that is the same for all panels). Initially, a pulse of FGF has little effect but pulses at intermediate time can have large effects. Time is normalized to be between 0 and 10. FGF feedback is not modeled here (see Figure 5A). (E) Sensitivity to timed FGF perturbations in the dual cusp given a constant bias towards Epi. An initial pulse is strong enough to give a large fraction of PrE but later pulses have no effect.

### 2.4 The geometrical model fits existing experimental data

We fit only the heteroclinic flip geometry to data since it does not require the parameter tuning of the dual cusp, though we are not aware of any data that explicitly excludes the dual cusp. Under the flip geometry, the dual positive state, NANOG+ (Epi) and GATA6+ (PrE) signifying ICM, begins as a shallow attractor but after it destabilizes the mutual antagonism between these two states occurs when they fall off the ridge separating the Epi and PrE. (The valley around the unstable ICM persists for some time).

Each cell sees the same potential landscape but evolves with an independent realization of the molecular noise term. There is a common but time dependent level of FGF as already justified in Figure 1. To complete the model, we have to define how the cells produce FGF. Previous models have considered differing molecular mechanisms for how the FGF is controlled by these two transcription factors. Ref. [15] considered FGF to be activated by NANOG whereas Ref. [13] considered it to be inhibited by GATA6. By modeling fates only, the geometric model allows us to be agnostic about whether the PrE state represses constitutive FGF expression or the Epi state is solely responsible for its production, or both. Similar agnosticism applies to the involvement of other transcription factors that reinforce the lineage decision between Epi and PrE.

In our flow diagrams, the proximity to the two fates is given by the *x* coordinate, +1 is pure Epi, and -1 pure PrE. Experiments indicate that cells which have specified into Epi can trigger the specification of undetermined cells into PrE but the reverse process does not take place [21]. We thus model FGF as proportional to the sum of the *x* coordinates of all cells, ignoring cells for which *x <* 0, i.e. any cell near the Epi state produces FGF. A linear dependence on *x* fits existing data, so nothing more elaborate is called for, though some experiments suggest a lag between the action of FGF and the cell state that could easily be accommodated. The effects of this feedback are shown in Figure 5A where we examine the effects of a pulse of FGF at different times. Early and late pulses have little effect (leading to WT difference between PrE and Epi) whereas intermediate pulses are most effective in changing fate.

**Figure 5:**
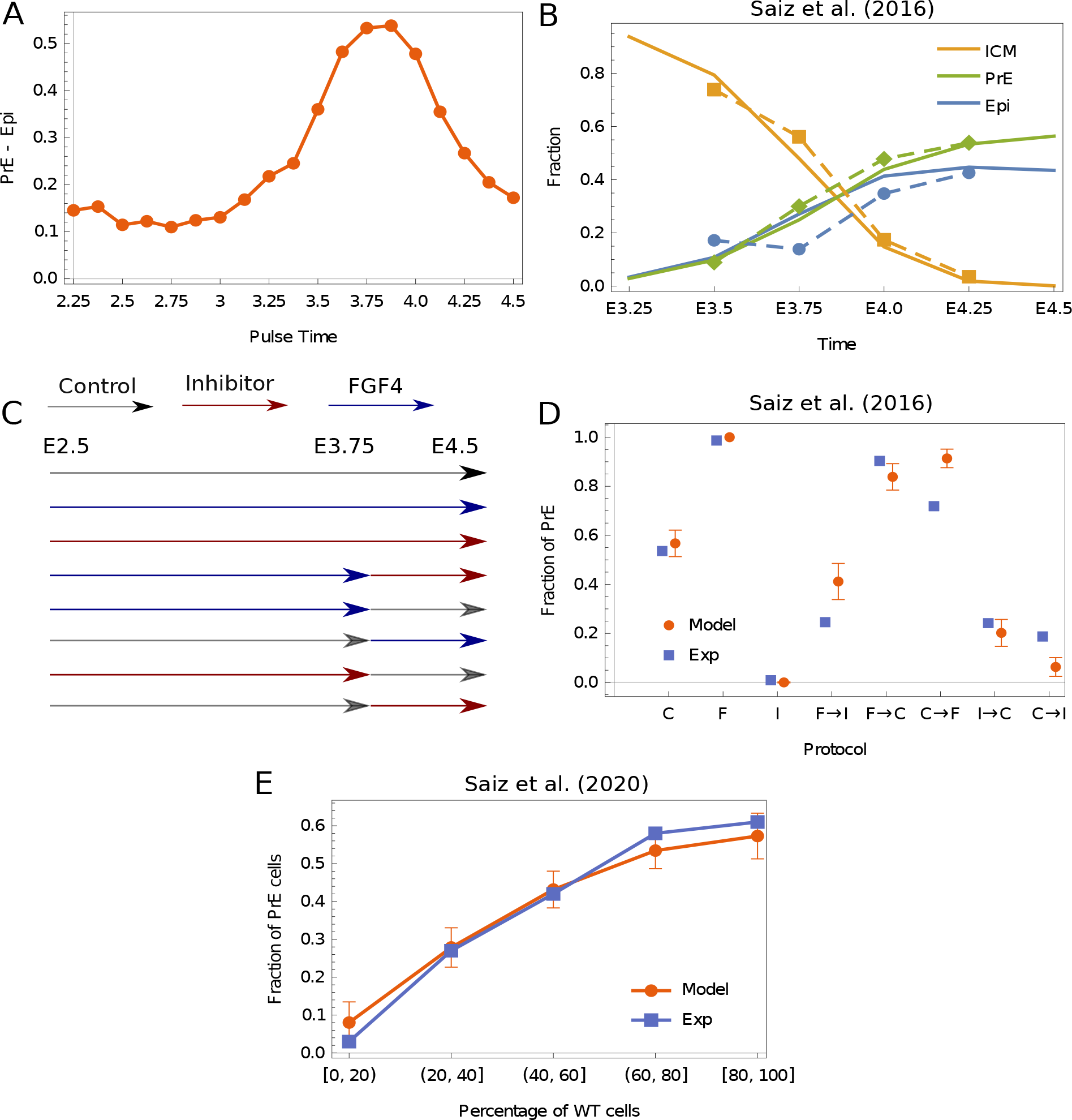
Theoretical model compared with the results of experiments. (A) Our model predictions with FGF feedback, for the effects of a pulse of FGF (magnitude 0.2, duration 0.5) plotted as the fraction difference of PrE and Epi. (B) Experiments from Saiz et al. [21] estimated the number of ICM/Epi/PrE cells by measuring NANOG and GATA6 expression at different times during the period of specification (dashed lines). Here, we show the analogous results from our model (solid lines). ICM cells were defined as those cells above a certain threshold in the *y* coordinate set equal to 0.5. Below this, cells were defined either as Epi *x >* 0 or PrE *x <* 0. We map embryonic time between E2.25 and E4.75 linearly to the time in the model. The bifurcation of the ICM state happens around E3 in the model. (C) Experimental protocols used by Saiz et al. [21]; embryos were kept in control media, media with inhibitor, and FGF4 rich media for the length of times indicated. (D) The results of our model for the perturbations in (C) compared with the experimentally measured results. The horizontal axes is marked by the protocol used with F standing for FGF4, C for control and I for Inhibitor. The model is qualitatively consistent with the data. The media was assumed to be switched exactly at E3.75 and a small uncertainty in the time when media is switched in can easily account for discrepancies. The error bars the theory result from stochastic fluctuations in the model. (E) The results of our model compared with experiments done by Saiz et al. [15]. Gata6^−*/*−^ cells were combined with wild-type cells in different proportions. The horizontal axis is the percentage of WT cells (which was grouped in different brackets). The vertical axis shows the fraction of PrE cells in the terminal state. This data sets a lower bound on the internal feedback required in the model.

To model a competence period, we allow the second parameter of our potential *K*_2_, which we have so far held fixed, to linearly decrease in time and transition through the value where the ICM state bifurcates away. (the molecular control of this decision is unknown, but the phenomena is very clear.) We map the computation time linearly between E2.25 and E4.75 corresponding to the point when most cells in the model are close to *x* = *±*1 i.e. Epi or PrE. The rate that *K*_2_ decreases is fit to the disappearance of ICM in Figure 5B.

The first experiments to define the competence window for response to FGF cultured the blastocyst in control media, and then added FGF4 or inhibited the MAPK pathway (arguably the converse) [14]. The specification process was captured in more detail by Saiz et al [21] by imposing thresholds on GATA6 and NANOG to define three states (which required us in the model to define a range of *y* above 0.5 corresponding to ICM). The transition from ICM to PrE or Epi occurred in an asynchronous fashion Figure 5B, that we fit. Dynamic noise leads to the error bars in Figure 5D.

Experiments in Ref. [22] divided the critical period for response to FGF4 into two hour intervals and saw suggestions of the temporal sensitivity we predict in Figure 4. However their protocol extended the exposure times in two hour increments rather than keeping the duration fixed and varying the application time. The maximal sensitivity of the cells to FGF4 application fell between E3.5 and E3.75 while FGF inhibition had it biggest effect between E3.5 and E4.5. As the authors note, this could be explained by a time lag in the production of FGF given the cellular state.

Finally recent experiments showed very directly that the blastocyst makes a population-level decision to control the ratio of Epi and PrE cells [15]. The authors combined *Gata6* null cells which are unable to specify PrE with WT in different proportions, and observed the fate of the WT cells. When the fraction of WT was below 60 % (the WT fraction of PrE) essentially all the WT cells converted to PrE cells indicating that internal population-level controls are very strong. We model the Gata6^−*/*−^ mutants by changing the initial conditions so they are already in the Epi state, and find essentially the same linear dependence of PrE fraction on WT between 0 and 60-80% WT (Figure 5E).

## 3 Discussion

### 3.1 A geometrical understanding of cell fate decisions

Previous models of the cell fate in the ICM have usually focused on the NANOG-GATA6-FGF/ERK regulatory network. The connections between these components are still not fully clarified and different models assume different kinds of regulation. It is difficult to precisely determine the parameters in such nonlinear models. Geometrical models bypass this difficulty by only modeling the cell fate. Furthermore when any gene centric model makes a decision between three fates, it must follow one of the bifurcations delimited by mathematics, so those bifurcations should be the subject of modeling from the start (the parameters in more elaborate models must collapse near the transition into those describing the bifurcation, and how they combine is problem dependent and was successfully negotiated in [17]).

Multiple genes define a lineage, and theory predicts that the dynamics of these genes are ultimately determined by the variables describing the bifurcation (as can be shown by taking a gene centric model and extracting the bifurcation). However, this determination can vary quantitatively as genes or combinations will relax at variable rates to the bifurcation pictures, so an underlying model to define the tendencies is essential for analyzing data. From the data in [11] the fate of individual cells given a 2hr sample of the ERK is not very predictive of the ultimate fate, but the embryo as a whole achieves an accurate partitioning of cells into Epi and PrE, evidently by averaging many sloppy decisions.

From a list of the allowed bifurcations, two in the present context (dual cusp and flip), it becomes easier to design experiments that distinguish them. Though we believe most labs would favor the flip when it’s defined, no experiment explicitly rules out the dual cusp, to our knowledge. Time dependent perturbations are an ideal way to distinguish models and a geometric visualization make their outcomes transparent, as illustrated by the qualitatively different responses of the dual cusp and flip to a pulse of FGF shown in Figure 4D,E. Similarly Figure 3, shows if a pulse of FGF is applied prior to lineage commitment the dual cusp requires a transition back through the ICM, while the flip does not. With a model, quantitative experiments that use a modest dose of morphogen to perturb outcomes are more informative than treatments that give all or none responses because they probe the boundaries between states.

Prior models of the ICM transition illustrate the difficulties incurred by being too specific. A model of Tosenberger et al. [13] has 4 dynamical equations with 24 parameters (plus additional equations for the FGF) but near the point of ICM instability, their model is equivalent to a dual cusp (Ref. [23] Figure S4 in their supplement). We have two equations with two free landscape parameters (one time dependent) in need of fitting (plus additional constants for the FGF). The model in Ref. [15] assumed only bistability between PrE and Epi in 1D, but had to initialize the cells at the dual positive (NANOG + GATA6) saddle point, which is difficult to imagine, though their data is consistent with the 2D flip geometry. The model in Ref. [16] proposed an inverted pitchfork bifurcation in which the ICM disappears to produce two new fixed points corresponding to Epi and PrE. The authors fixed the cells to a 2D lattice and assumed variable range FGF signaling, so the spatial geometry was tied to the cell fates, in a manner similar to Turing systems with boundaries or Notch-Delta repression. The data in Ref. [11] shows considerable cell movement and PrE markers that appear before spatial segregation, so we consider the conclusions in [16] to be too contingent on assumptions.

We have emphasized the flip topography for its logical consistency (i.e., we avoid the parameter tuning inherent in the dual cusp, or pitchfork). The flip, being a global bifurcation, is ideally suited to transitions where the embryo needs to control the population ratio via morphogen feedbacks. The transition involves a reconnection of the orbit leaving the progenitor from one terminal state to the other. In our case, the progenitor destabilizes but in other situations, it remains an attractor and cells dribble out via fluctuations [20]. Signals have a finite time to act on the orbit and effect the flip.

### 3.2 Model Assumptions and their Validity

A clear difference between our model and previous models in the literature is the role that spatial interactions play. Most previous models have considered the FGF signaling to mediate very localized spatial interactions. A recent study measured the spatial range of FGF signaling in a mouse Embryonic Stem-Cell System by creating a cell line with an FGF4 transcriptional reporter [24]. They estimated that the spatial range of FGF signaling was 2 spatial neighbors.

While it may be true that FGF signaling *in vivo* is also somewhat localized, we do not find evidence of spatial correlations at the level of the ERK activity, perhaps due to cell movement. In our model, we assume a single homogeneous but time dependent FGF concentration. Current data does not force us to contemplate anything more complex.

However Ref. [5] infers the need for localized FGF interactions working with blastocysts with *Fgf4* −*/*− knocked out. They found that exogenous FGF applied during the growth from 32-100 cells gave embryos that were either entirely Epi or entirely PrE at the end, but passed through an earlier stage of mixed NANOGGATA6 expression. Exogenous FGF did not rescue the knockout, yet the FGF level was not extreme as evidenced by the dual outcomes (all Epi or all PrE). Perhaps FGF needs to be finely tuned to get a balanced outcome in the absence of feedback. It would be interesting to repeat this experiment with a live ERK reporter and track cell motility, to see if the dynamics replicates Ref. [11]. It may also point to a backup mechanism for enforcing homogeneous fates based on integrin or cadherin mediated contacts

We have assumed that one parameter in the landscape controls the bifurcation of the ICM and a separate parameter receives FGF input. In reality the two are probably mixed, but current experiments do not allow us to disentangle the mixing. The model relating the internally generated FGF to the cell state, is phenomenological. We have assumed the readout of cell state is instantaneous, but time lags are plausible.

Cell division and the correlation between the fates of the daughters plays no role in our model. If a cell divides in a diverging region of the flow plane, the correlation between the daughters would tend to make a final Epi to PrE ratio a bit noisier than in the absence of correlation, but that statistic does not yet figure in the model fits. The effect could be mimicked by adjusting our noise source.

Our competence period of the cells for the FGF signal is set by a separate event that bifurcates the ICM at a specified time. Such signals are known in other contexts. For example, in germ layer specification in zebrafish, a recent study reported that Nodal signaling set the competence window for FGF to regulate fate [25]. Our assumption is consistent with previous studies that show the cells becoming competent to FGF signaling at a specified time post fertilization [26]. How the embryo keeps time or counts cells is an open and interesting question.

In addition to timing, experiments where the levels of FGF and inhibitors are titrated down towards threshold would define a presumably sigmoid response to FGF (and by comparison, the level for what is produced internally). Partial penetrance is a way to infer the boundaries between fates not a hindrance as it would be in genetic screens. Experiments that control timing and deal with partial penetrance require a model since the outcomes are quantitative not binary. Ultimately, our model seeks to encourage and help interpret future experiments such as we propose in Figure 4 to study this paradigmatic transition in greater detail.

## Acknowledgments

AR acknowledges support from the Department of Atomic Energy, India under project No. RTI4006, and the Simons Foundation Grant No. 287975. EDS was supported in part by (US) National Science Foundation Grant PHY-1748958. AR was a fellow at the Rockefeller Center of Studies in Physics and Biology where this project was initiated. J. Briscoe, J. Delas, A.K. Hadjantonakis, W. Hur, and E. Tanaka made helpful comments on an earlier draft of this paper. Nestor Saiz and Claire Simon provided additional insights to the experiments they did in the Hadjantonakis lab.

## 4 Materials and Methods

### Spatial Correlations and Lineage Statistics

The correlations in Figure 1 are calculated for the data in Ref. [11]. The spatial correlations are calculated using the 2 hr movies (imaged at 5 minute intervals). We remove all cells which have the TE fate (manually labeled). We estimate the average time taken for one neighbor for a cell to change by taking every embryo, calculating the Voronoi neighbors for each cell, and calculating the average time it takes for one unique Voronoi neighbor to change (i.e. if the same neighbor is lost and gained in the 2 hour time period, we count it only once). This turns out to be around 25 minutes in the data. We thus measure spatial correlations from data separated by 30 minutes (i.e. 4 data points from one 2 hour movie). We measure spatial correlations using the Moran’s I index. We use a weight matrix *w*_*ij*_ which is row-standardized so Σ_*j*_ *w*_*ij*_ = 1. The statistic is then given by

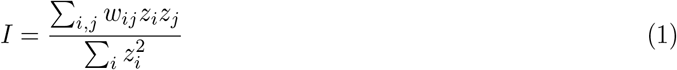

where *z*_*i*_ is the data which is centered around the mean. Given a physical distance *d*_*ij*_ between two cells indexed by *i* and *j*, we define *w*_*ij*_ = *H*(*d*_*ij*_ − *d*)*/Σ* _*j*_ *H*(*d*_*ij*_ − *d*) where *H* is the Heaviside theta function which is 1 for distances below some fixed distance *d* = 20 *μ*m and 0 otherwise. The value of *d* = 20 *μ*m is approximately the mean distance between Voronoi neighbors in the data. We calculate this statistic for snapshots of embryos.

The lineage statistics are calculated from the 12 hr movies (imaged at 15 minute intervals). We take all cells which have undergone a mitotic division, and plot their mean ERK C:N Ratio as well as the absolute difference between the ERK C:N Ratio for the two sisters for a 2 hour time period (8 data points).

### Equations for Potentials

The potentials for the dual cusp and heteroclinic flip were modified from Ref. [20]. For the dual cusp

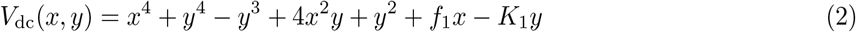

For the heteroclinic flip

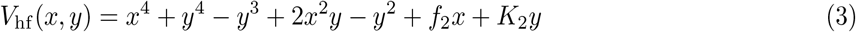

For Figure 2, we used *K*_1_ = 0.15, *K*_2_ = 1.4, *f*_1_ = 0, *f*_2_ = 0. The potentials in Figure 2 are schematic only drawn to emphasize the topology. The flow lines and bifurcation diagram are drawn using the above equations.

For a given potential *V* (*x, y*), the flow is modeled as

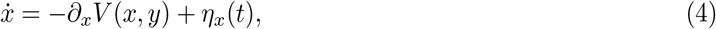

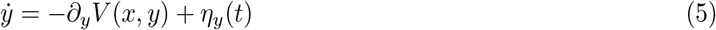

where *η*_*x*_(*t*) and *η*_*y*_(*t*) are independent noise terms such that ⟨*η*_*x*_(*t*)*η*_*x*_(*t*^*’*^) ⟨ = *σ*^2^*δ*(*t* − *t*^*’*^) and ⟨*η*_*y*_(*t*)*η*_*y*_(*t*^*’*^) ⟨ = *σ*^2^*δ*(*t*−*t*^*’*^). We choose *σ* = 0.05. The initial condition is chosen as a small Gaussian Distribution (Standard Deviation 0.05) around the fixed point corresponding to the ICM state for zero FGF concentration and *K*_1_ *≈* 1.45 (*x* = 0, *y* = 0.8) for the flip, and *K*_2_ *≈* 0 (*x* = 0, *y* = 0) for the dual cusp.

For Figure 3, we chose *K*_1_ = − 0.05 and *K*_2_ = 1.55. The initial conditions for the dual cusp are drawn from a normal distribution with mean 0.3 and −0.2 in *x, y*, which is towards the Epi state. For the heteroclinic flip, the initial conditions are drawn from a normal distribution with mean 0.1 and 0 respectively, towards the Epi state. In both cases, *σ* = 0.05 and external FGF *f*_1_, *f*_2_ = 0.2. Streamlines are shown at the bifurcation of the ICM state.

For Figure 4, we chose *K*_1_ = −0.05 and *K*_2_ = 1.55, *σ* = 0.05. The external FGF is given as a pulse of magnitude 0.2 and duration 0.5. Identical realizations of a slightly larger noise (*σ* = 0.1) are used in A-C for demonstrative purposes. Streamlines are shown at the bifurcation of the ICM state. The plots in D-E are averages over 2500 runs of single cells.

### Dynamical equations for Full Model

For the full model cells interact via the common FGF and follow the flip landscape. Each one of them has the equation

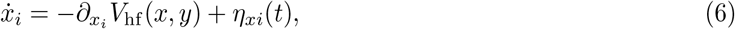

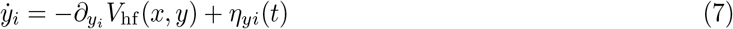

The noise terms are defined as before with each cell seeing a different realization sampled with an identical standard deviation *σ* in both directions. The time of cell fate specification is normalized to be between 0 and 10 with a scale factor to embryonic time *τ*. Time *t* = 10 is assumed to be the end of the competence period when *f*_1,2_ revert to 0. The bifurcation of the ICM state is done globally for each of the cells by making *K*_2_ = *m*_0_ + *m*_1_*t* with *m*_0_ = 1.45 and *m*_1_ = 0.022, where *t* is time. The bifurcation happens around *t ≈* 2.5. The parameter *m*_1_ controls how rapidly cells leave the vestige of the ICM state. (In [20] the analogue of the ICM state never lost stability and the cells exited due to the noise.) Since data is available as in Figure 5, this allows us to fix our map with *t* = 0 equivalent to E2.25 and *t* = 10 equivalent to 4.75 and a unit difference corresponding to 0.25 days.

The FGF concentration

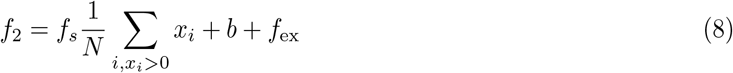

where the first contribution is only from those *x*_*i*_ that are positive and represents the feedback from the Epi cells. The second is an initial bias towards the Epi state and the third is the effect of external FGF. We set *b* = −0.004, *f*_*s*_ = 0.1, *f*_ex_ = 0.2 for FGF over-expression and −0.3 for the inhibitor. In principle, the inhibitor would affect the *f*_*s*_ term which we have not modeled. *N* is the number of cells (taken to be 50).

### Comparison with Experiments

The coefficients in the polynomials in Eq. 2,3 other than those denoted symbolically are not fitting parameters. Other choices that differ modestly from those shown could be absorbed by a change in variables. Since we associate fates with fixed points and the topology of the flows connecting them will not change, the variable change is of no consequence.

We have 7 important parameters in our model: three to define the FGF feedback in Eq.[8], three for the time scales (the slope and origin of *K*_2_, and the mapping of embryonic to computational time), and one to delimit ICM for the terminal fates (Figure 5A). Of these seven, three are unavoidable definitions of units (of FGF concentration and time). The threshold to delimit ICM is unavoidable. The bias in FGF is needed to capture the asymmetry in PrE and Epi. The form of *K*_2_ is required to fit the competence period and asynchronous loss of ICM character. Apart from these seven, the noise in the equations *σ* and the variability in timing of the FGF signal is less important. The initial conditions around the ICM state are similarly not very important.

We set the time scales by comparing to Figure 5 (B). We set the scale of the internal FGF by comparing to Fig 5(E). The scale of the external FGF is set by comparing to Figure 5D. The media is assumed to be switched exactly at E3.75 but it is possible to assume a slight variability in time which could explain the discrepancies from data. The data is taken from Ref. [21] Figure 1-h, Figure 5-f and Figure 2-d.

For comparing to the experiments shown in Figure 5(E), a random proportion of cells are assumed to be in the Epi state (the Gata6 mutants). The rest are assumed to start at the ICM state. We then run our simulations several times (1000 runs) and bracket our results the same way as in Ref. [15] by the number of cells that are mutants vs wild-type. The data is taken from Figure 1(S2)-L in Ref. [15].

